## Supplemental information for "Molecular chaperone BiP controls activity of the ER stress sensor Ire1 through interactions with its oligomers"

**List of Abbreviations**

ATF6 – activating transcription factor 6

BiP – Immunoglobulin binding protein

CD – Cytoplasmic domain

DLS – Dynamic light scattering

DTT – Dithiothreitol

E. coli – Escherichia coli

EDTA – Ethylenediaminetetraacetic acid

EM – Electron microscopy

ER – Endoplasmic reticulum

FIDA – Flow-induced dispersion analysis

FITC – Fluorescein isothiocyanate

HEPES – 4-(2-hydroxyethyl)-1-piperazineethanesulfonic acid

HMK – 20 mM HEPES, 100 mM KCl, 5 mM MgCl_2_ buffer

HMQC – Heteronuclear single quantum coherence

HSP – Heat shock protein

INEPT – Insensitive nuclei enhancement by polarization transfer

IPTG – Isopropyl β-D-1-thiogalactopyranoside

IRE1 – Inositol requiring enzyme 1

LB – Lysogeny broth

LD – Luminal domain

LIC – Ligation independent cloning

MS – Mass spectrometry

MST – Microscale thermophoresis

Mw – Molecular weight

NBD – Nucleotide binding domain

NMR – Nuclear magnetic resonance

OD – Optical density

PDB – Protein data bank

PERK – Protein kinase R-like endoplasmic reticulum kinase

PONDR – Prediction of natural disordered regions

PQC – Protein quality control

SDS-PAGE – Sodium dodecyl sulphate polyacrylamide gel electrophoresis

SEC – Size exclusion chromatography

TEV – Tobacco etch virus

TROSY – Transverse relaxation optimized spectroscopy

UPR – Unfolded protein response

UV – Ultraviolet

WT – Wild type

XBP1 – X-box binding protein 1

**Supplementary Figure 1: The oligomeric state of apo WT Ire1-LD and its D123P variant monitored by microscale thermophoresis (MST).**

MST traces were obtained from 0.5 µM FITC labelled Ire1-LD titrated with unlabelled Ire1-LD. Either WT Ire1-LD or the dimerization-deficient D123P variant^1^ was used for these measurements. Only WT Ire1-LD titration by WT Ire1-LD shows the dimerization pattern (black). As expected, the titration of D123P variant by either the D123P variant (blue) or WT Ire1-LD (red) results in no dimerization. Error bars indicate ±SE (standard error) for three replicate experiments.

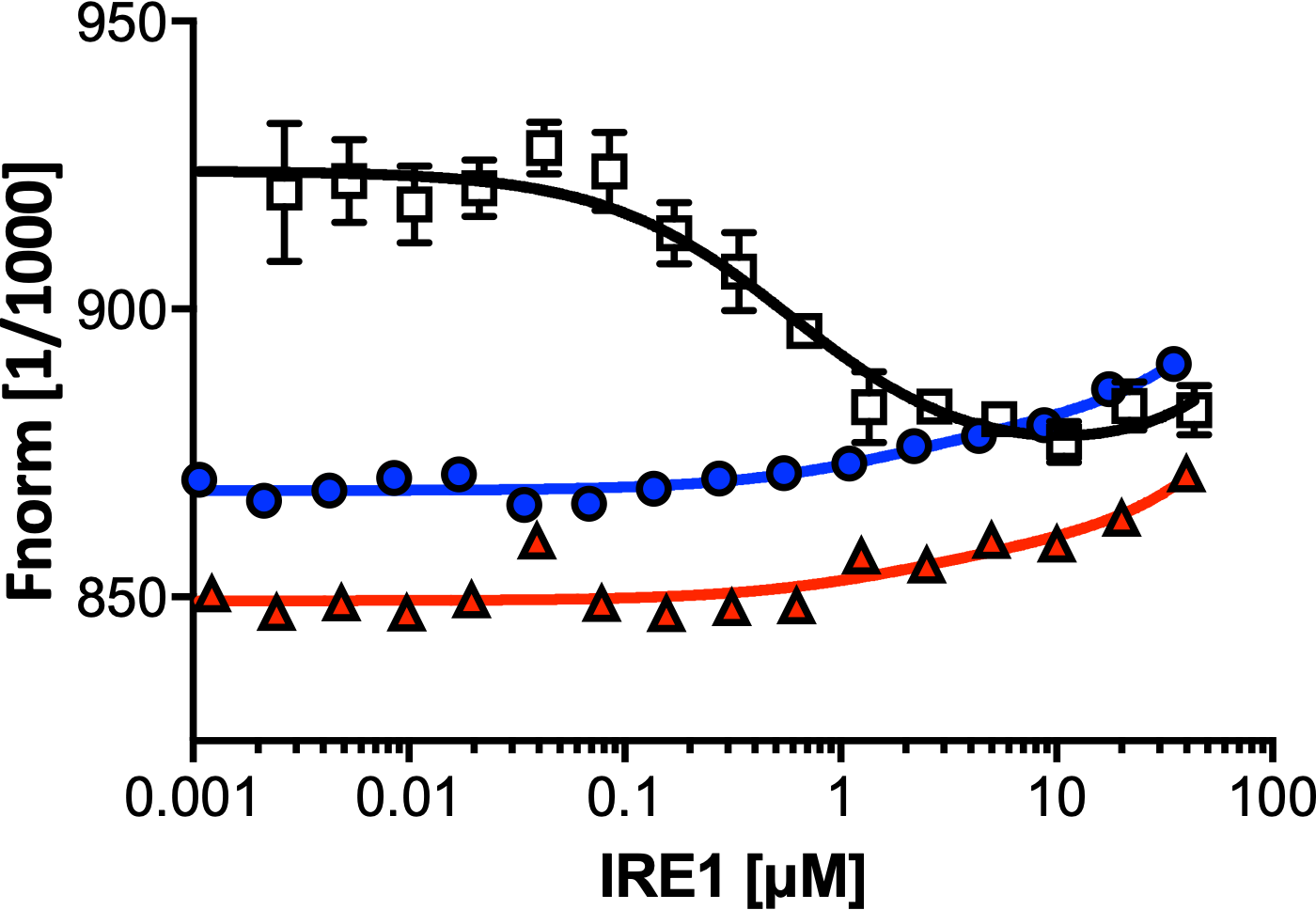

**_WT Ire1-LD* + WT Ire1-LD_**

**_D123P Ire1-LD* + D123P Ire1-LD_**

**_D123P Ire1-LD* + WT Ire1-LD_**

**Supplementary Figure 2: The oligomeric state of apo WT Ire1-LD and its D123P variant monitored by size exclusion chromatography (SEC).**

SEC elution profiles of WT Ire1-LD (a) and the dimerization-deficient D123P variant^1^ (b) at different protein concentrations (from 7.5 to 60 μM, coloured as annotated); the fraction of monomeric Ire1-LD for Fig. 1a was calculated from the peak position at the corresponding concentration of Ire1-LD. For calculations of peak positions, we assumed that the D123P variant is monomeric at these concentrations.

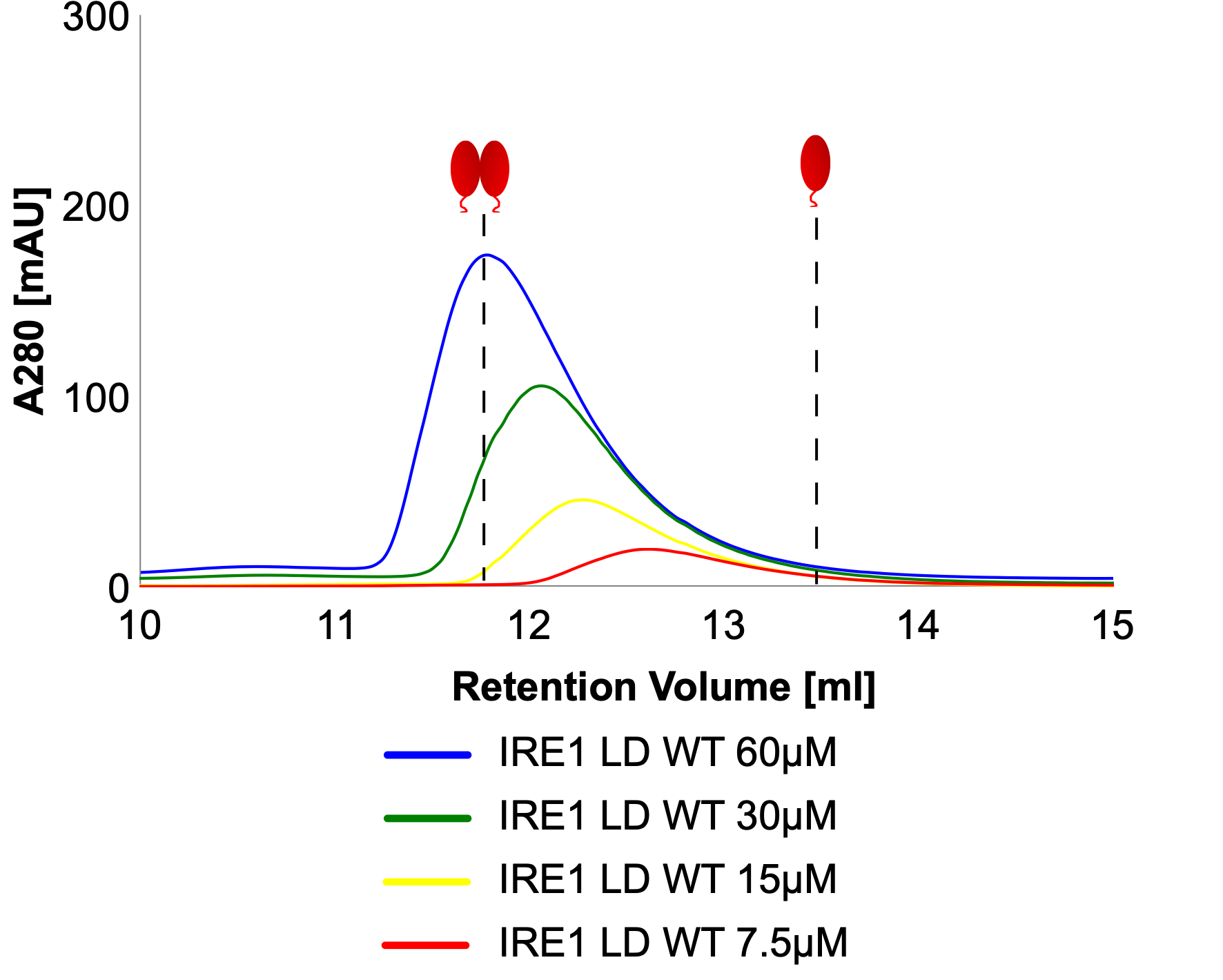

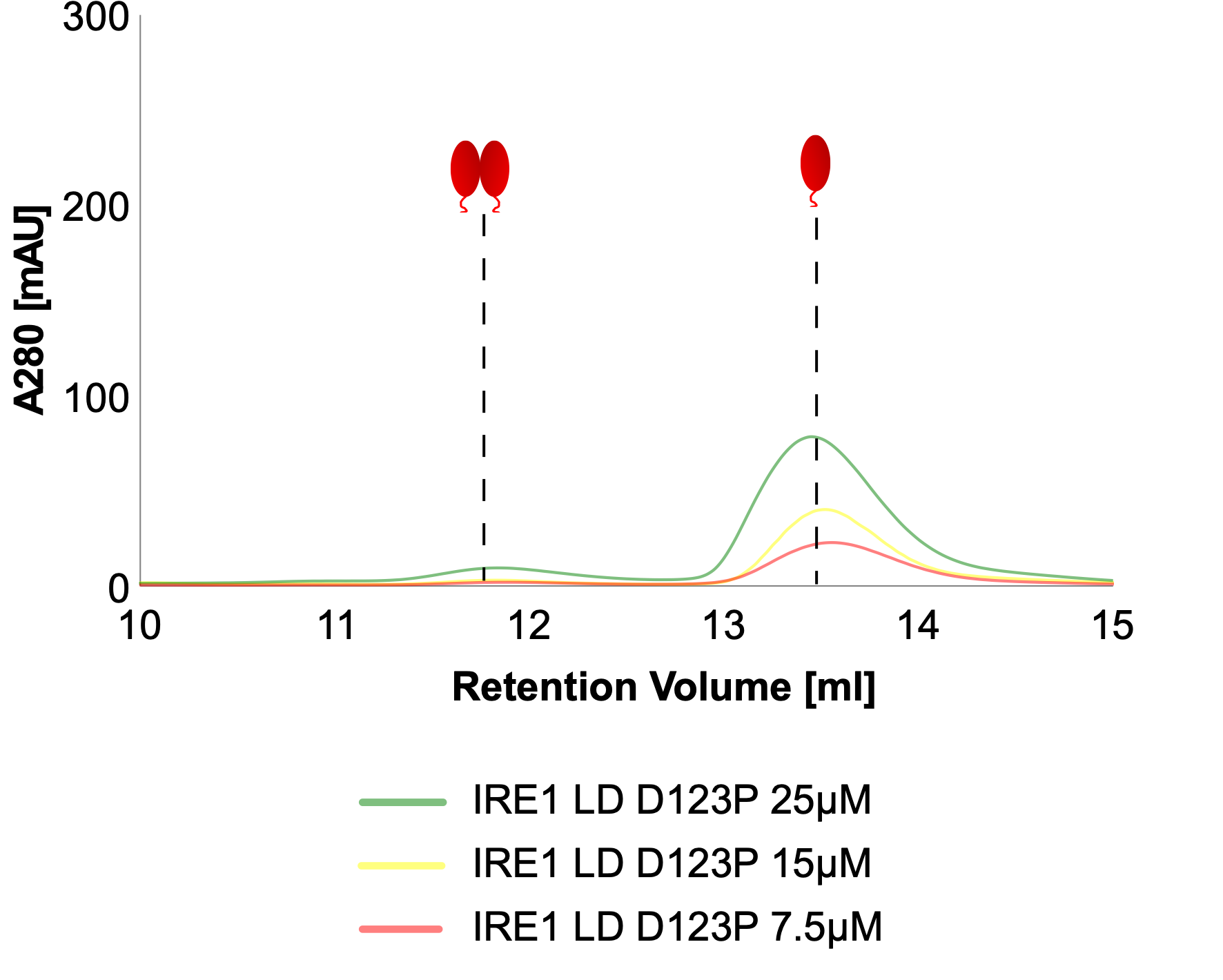

**a**

**b**

**Supplementary Figure 3: The oligomeric state of apo WT Ire1-LD monitored by native mass spectrometry.**

Native MS of 5 µM Ire1-LD showing the presence of monomers (highlighted in red, Mw is 49658.65+-1.17), dimers (highlighted in black, Mw is 99320+-1.68) and tetramers (highlighted in blue, Mw is 198796.66+-11.01). The sample was sprayed from 100 mM ammonium acetate at pH 6.8.

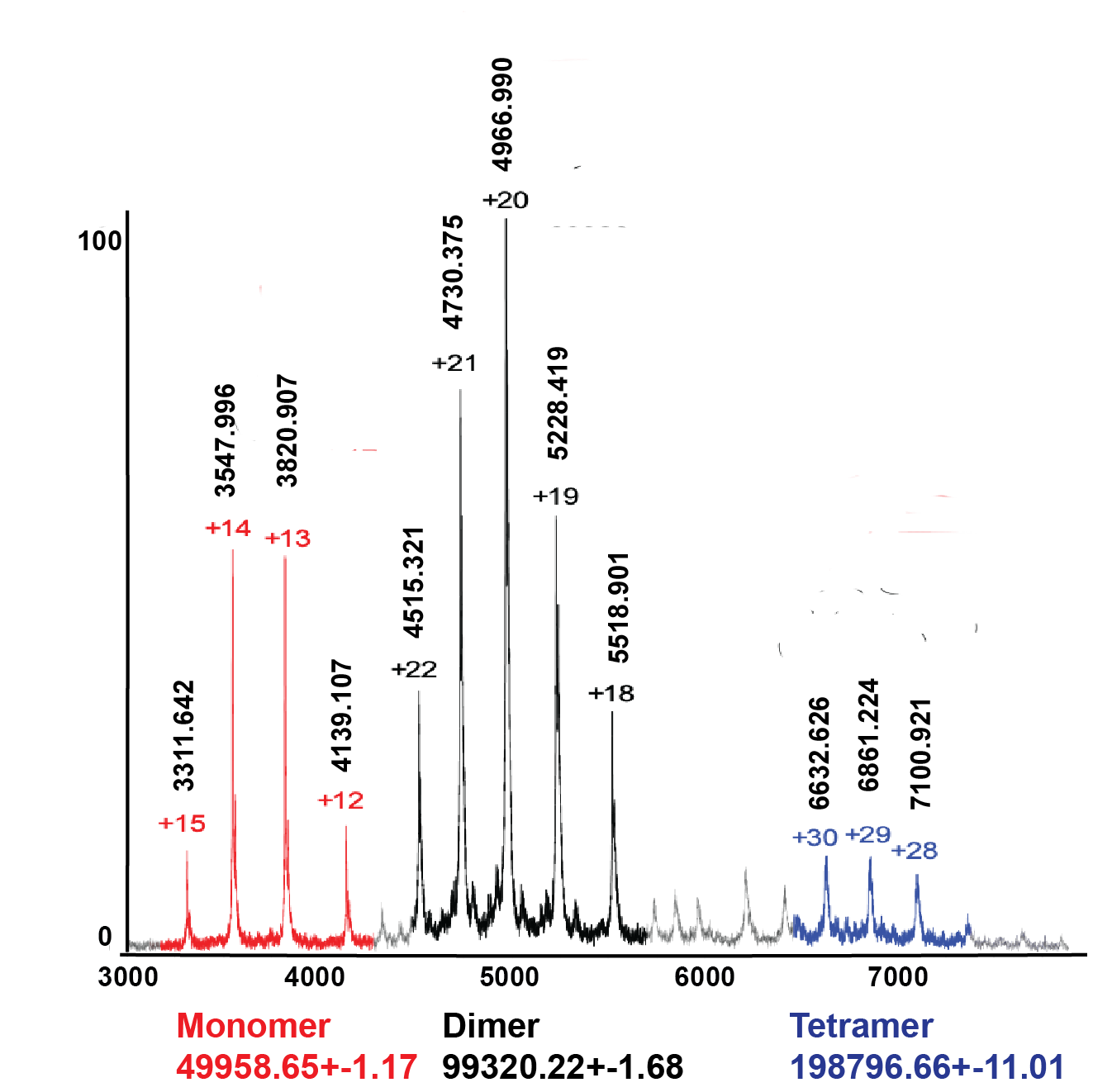

**% Intensity**

**m/z (Da)**

**Supplementary Figure 4: Formation of insoluble Ire1-LD oligomers in the presence of the high-affinity ΔEspP substrate.**

(a) The turbidity measurements of 20 µM Ire1-LD in the presence of ΔEspP at different (0-128 µM) concentrations. Error bars represent standard deviations for replicate experiments.

(b) The amount of insoluble Ire1-LD as a function of ΔEspP measured by the solubility assay. The amount of insoluble Ire1-LD was calculated as the total Ire1-LD concentration (20 µM) minus the amount of soluble Ire1-LD measured by the Bradford assay.

(c) The calibration curve^2^ of the amount of insoluble Ire1-LD oligomers vs. OD at 400 nm for the corresponding ΔEspP concentration.

(d) The amount of aggregated protein over time calculated from (a) using the calibration curve from (c) as a function of time. The fraction of insoluble Ire1-LD oligomers as a function of ΔEspP concentration was calculated as described by Borgia et al.^2^ using the turbidity measurements at different ΔEspP concentrations (a) and the amount of insoluble protein (b) as a function of ΔEspP.

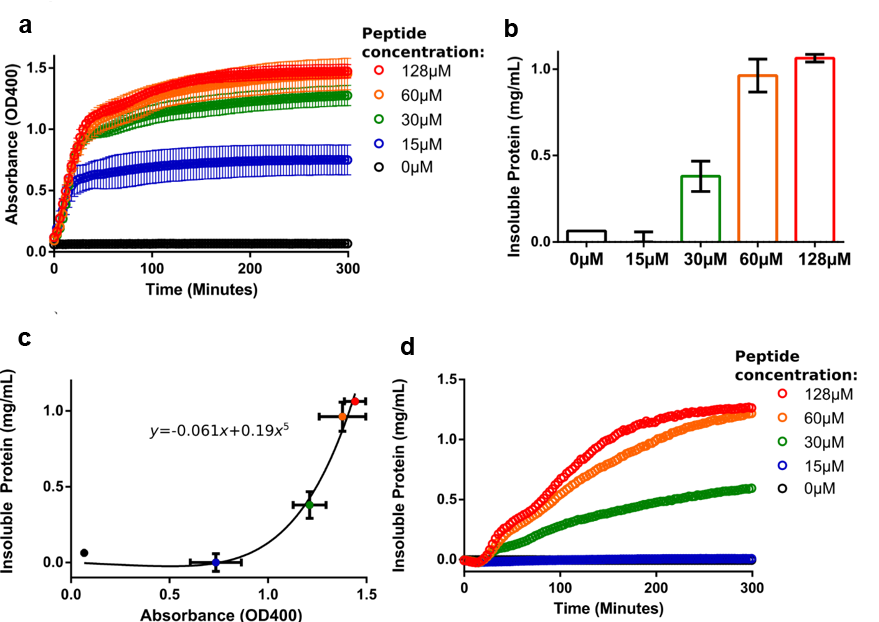

**Supplementary Figure 5**: **Substrate-induced oligomerisation of Ire1-LD monitored by fluorescence polarization.**

(a) The fluorescence polarisation assay to measure ΔEspP-dependent oligomerisation of Ire1-LD. 50 nM FITC-labelled Ire1-LD was incubated with increasing concentrations of ΔEspP for 30 minutes; the apparent constant for this peptide-dependent oligomerisation is 20.6±1.13 µM. Error bars indicate ±SE (standard error) for 3 replicate experiments.

(b) The fraction of insoluble Ire1-LD oligomers obtained from the turbidity measurements (Supplementary Figure 4d). Error bars indicate ±SE (standard error) for three replicate experiments. The apparent constant for this ΔEspP-dependent formation of insoluble oligomers is 30.5±5 µM, in good agreement with K_1/2_ of formation of soluble oligomers obtained from fluorescence polarisation assay (a).

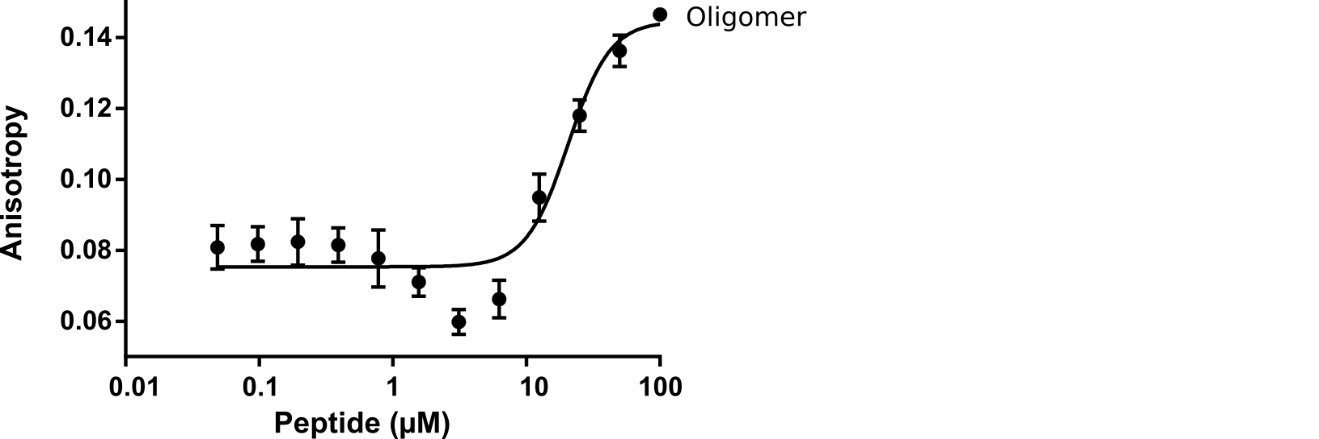

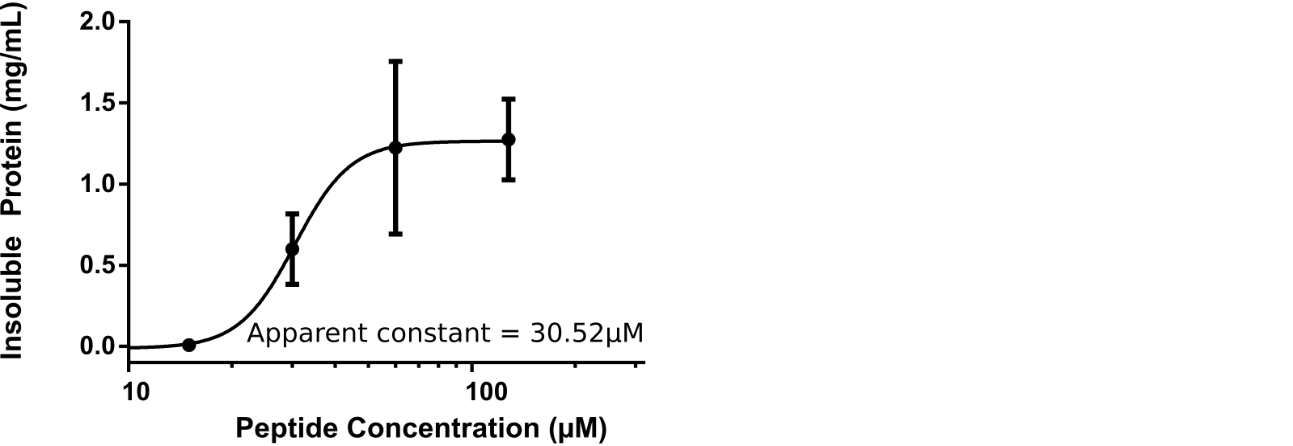

**a**

**b**

**Supplementary Figure 6: Formation of insoluble Ire1-LD oligomers in the presence of the low-affinity MPZ1 substrate.**

The turbidity measurements of 20 µM Ire1-LD in the presence of MPZ1 at different (0-359 µM, annotated) concentrations. Only the highest (359 μM) concentrations of MPZ1 promoted the formation of insoluble Ire1-LD oligomers. Error bars represent standard deviations for replicate experiments.

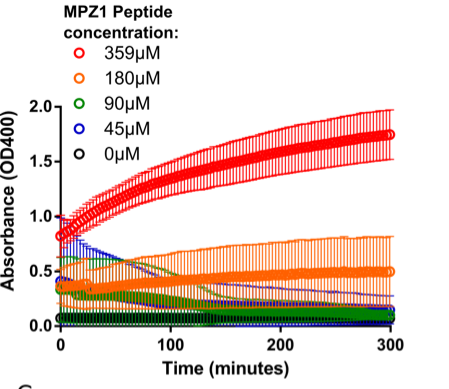

**Supplementary Figure 7: Perturbations in the dimerisation and oligomerisation interfaces significantly disturb the formation of insoluble Ire1-LD oligomers**

Turbidity measurements of D123P (a) and ^359^WLLI^362^ to GSSG (b) variants of Ire1-LD in the presence of ΔEspP at different concentrations. Error bars represent standard deviations for replicate experiments. (c) The fraction of insoluble Ire1-LD and its variants (calculated from turbidity assay as described in Supplementary Figure 4c,d) vs ΔEspP concentration. For both variants, the formation of insoluble oligomers was observed at significantly higher peptide concentrations as compared with the WT Ire1-LD, demonstrating that perturbation of Ire1-LD dimerization (in the D123P variant) and formation of soluble oligomers (in the ^358^WLLI^362^ to GSSG variant) significantly affect the formation of insoluble Ire1-LD oligomers.

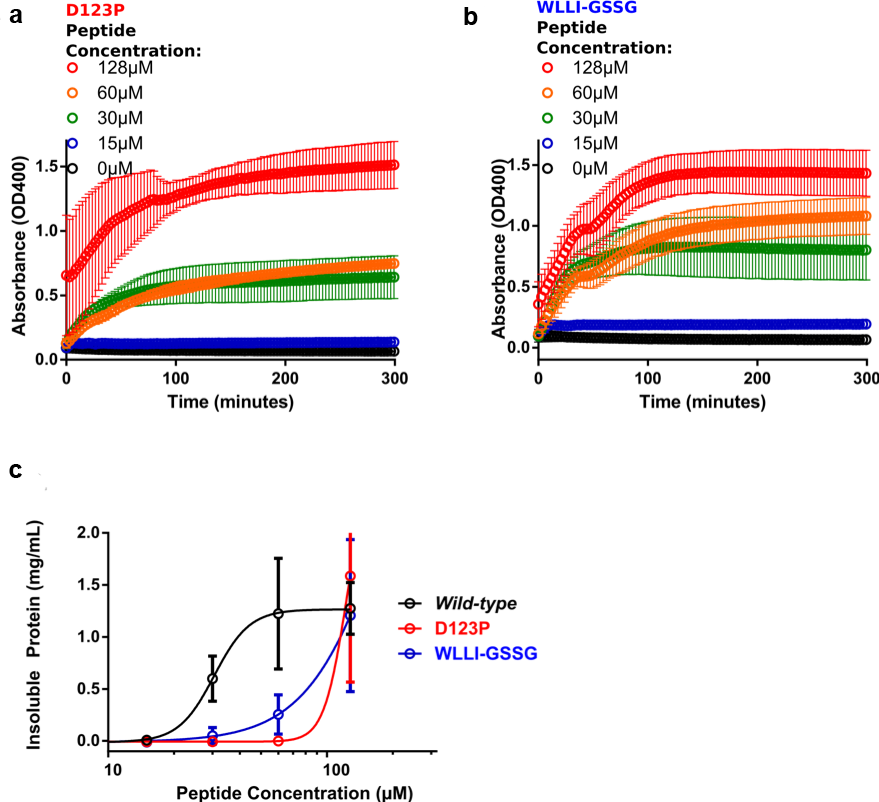

**Supplementary Figure 8: The building blocks of Ire1-LD insoluble oligomers are folded Ire1-LD dimers.**

(a) ^1^H-^15^N 2D correlation ssNMR spectra of insoluble Ire1-LD oligomers formed in the presence of the ΔEspP recorded using cross-polarisation (CP, red) and INEPT (black) based ^1^H-^15^N polarisation transfer. The INEPT-based spectrum (black) contains very few peaks, suggesting that the oligomers contain only few long-disordered regions with high mobility on the fast timescales and the most of the protein is immobilised; furthermore, the CP spectrum (red) has a good dispersion of ^1^H peaks, indicative of a folded protein conformation.

(b) A plausible model structure of Ire1-LD oligomers based on the Alphafold model of human Ire1α-LD (AF-A0A7P0TAB0, only residues 24 to 365 are shown); the model was built in PyMol using the dimerization and oligomerization interfaces previously suggested by ^1,3^.

(c) A representative TEM image of insoluble Ire1-LD oligomers formed in the presence of the ΔEspP peptide (same as in a), demonstrating that the diameter of the elongated oligomers is consistent with the diameter of the Ire1-LD dimer in the model structure (from b).

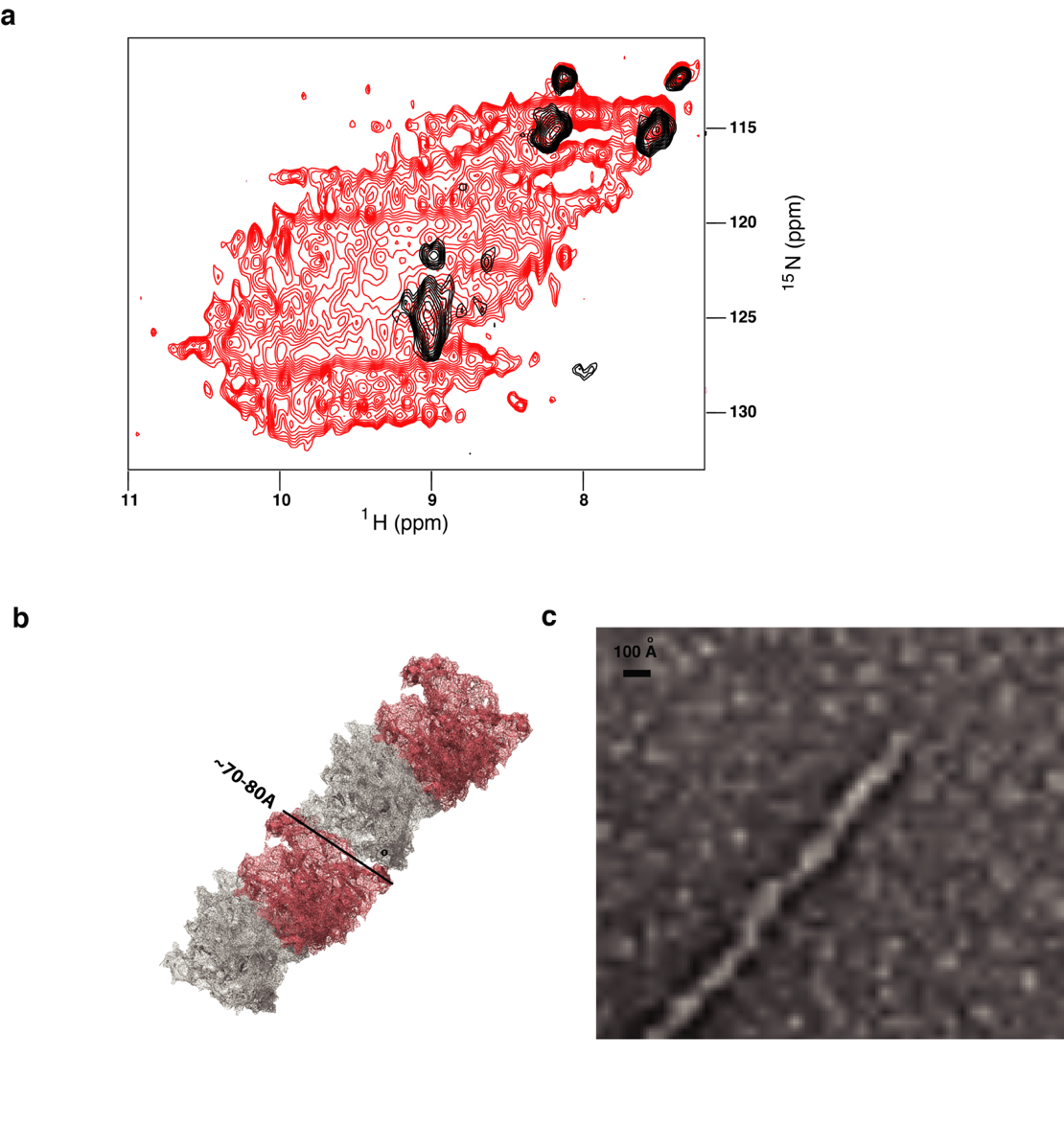

**Supplementary Figure 9:** **The size of Ire1-LD oligomers characterized by flow-induced dispersion analysis (FIDA)**.

(a) Representative Taylograms for samples containing 10 µM of Ire1-LD and 1 µM ΔEspP in the presence and absence of BiP and ATP. The 1:10 ΔEspP:Ire1-LD ratio was used to achieve a good signal-to-noise ratio but avoid the formation of insoluble oligomers during the experiments.

(b) The apparent hydrodynamic radius of Ire1-LD oligomers in the absence and presence of BiP. Sub-stochiometric concentrations of BiP [1:10 (1 µM BiP) and 1:100 (0.1 µM BiP) in red and pink, respectively] were sufficient to reduce the size of Ire1-LD oligomers. Error bars indicate ±SE (standard error) for at least three replicate experiments.

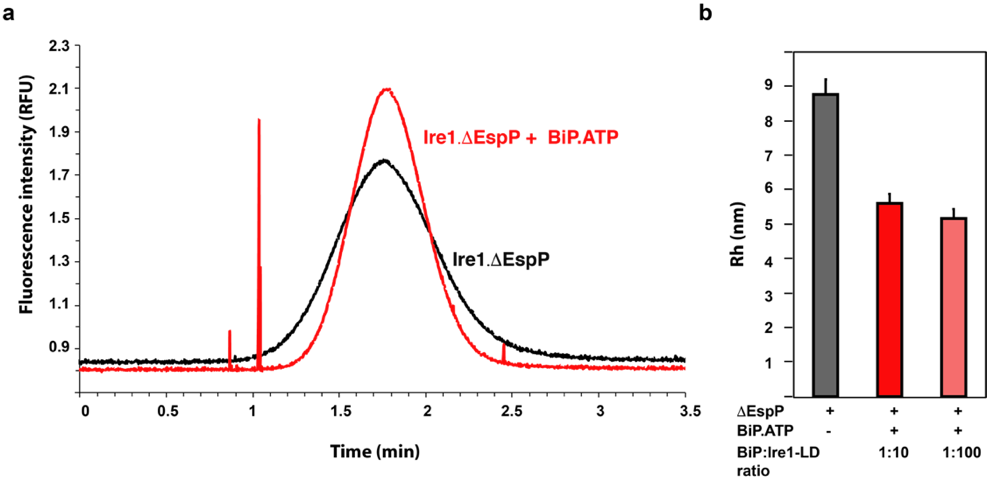

**Supplementary Figure 10: BiP does not form a stable complex with small Ire1-LD oligomers.**

The isoleucine region of methyl-TROSY spectra of ATP-/ADP-bound and apo full-length U{^2^H,^12^C}, Ile-C^δ1^-^13^CH_3_ WT BiP (in black) overlaid with the spectra of corresponding nucleotide-bound state of BiP in the presence of 50 μM Ire1-LD and 100 μM ΔEsp (in red). Ire1-LD was preincubated with BiP and if required 40 mM ATP for one hour and the spectra were recorded immediately after the addition of ΔEsp to monitor BiP interactions with NMR-visible Ire1-LD species (by monitoring changes in peak positions and peak intensities upon interactions). Due to ATP hydrolysis during the experiments, peaks for both ATP- and ADP-bound BiP are present. Within experimental time (ca. 30 min), no significant changes in peak intensities/positions were observed. The absence in any changes in the BiP spectra demonstrates that either ATP-bound or unbound BiP do not interact with either soluble Ire1 species or ΔEsp. In the absence of ATP, after incubations for longer than 1 hour, BiP peaks in NMR spectra gradually decreased due to BiP interactions with insoluble Ire1-LD.ΔEsp oligomers.

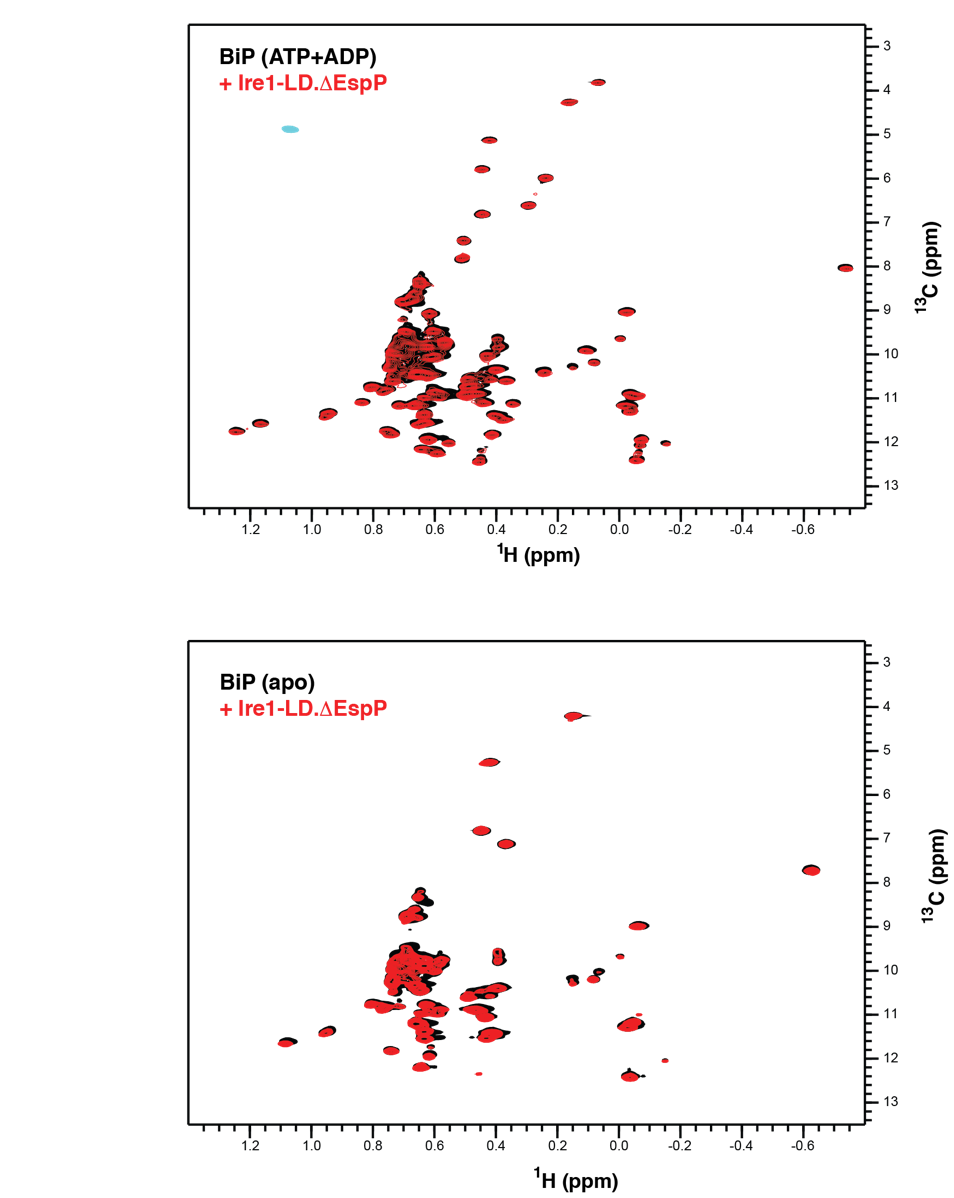

**Supplementary Figure 11: NMR identification of BiP binders.**

(a) The representative isoleucine region of methyl-TROSY spectra of ATP-bound full-length BiP* T229G in the presence of the model substrate HTFPAVL (1 mM) and Ire1-derived binders: 50 μM GSTLPLL, 50 μM RNYWLLI, or Ire1-derived non-binders: 1 mM AVVPRGS, 1 mM KHRENVI, 1 mM ENVIPADS or 1 mM KDMATIIL (Supplementary Table 1). The full spectra are shown in (c). Binding to the substrate (either HTFPAVL, GSTLPLL or RNYWLLI) stabilizes the domain-undocked conformation of BiP as monitored by a decrease in the intensities of peaks, corresponding the domain-docked conformation and an increase in intensities of peaks corresponding the domain-undocked conformation.^4^

(b) Bar chart showing the percentage of domain docked conformation for the three representative doublet peaks from (a), calculated as described previously^4^. Error bars indicate ±SE (standard error) for three doubles. The asterisk (*) respresents the p-value of the statistical test. ***p<0.0001.

(c) The full methyl-TROSY spectra of ATP-bound full-length BiP* T229G in the presence of the model substrate HTFPAVL (1 mM), Ire1-derived binders: 50 μM ^310^GSTLPLL^316^, 50 μM ^356^RNYWLLI^362^, or Ire1-derived non-binder: 1 mM ^388^ENVIPADS^395^ (the other three non-binders have identical spectra). The boxes highlight the regions shown in (a).

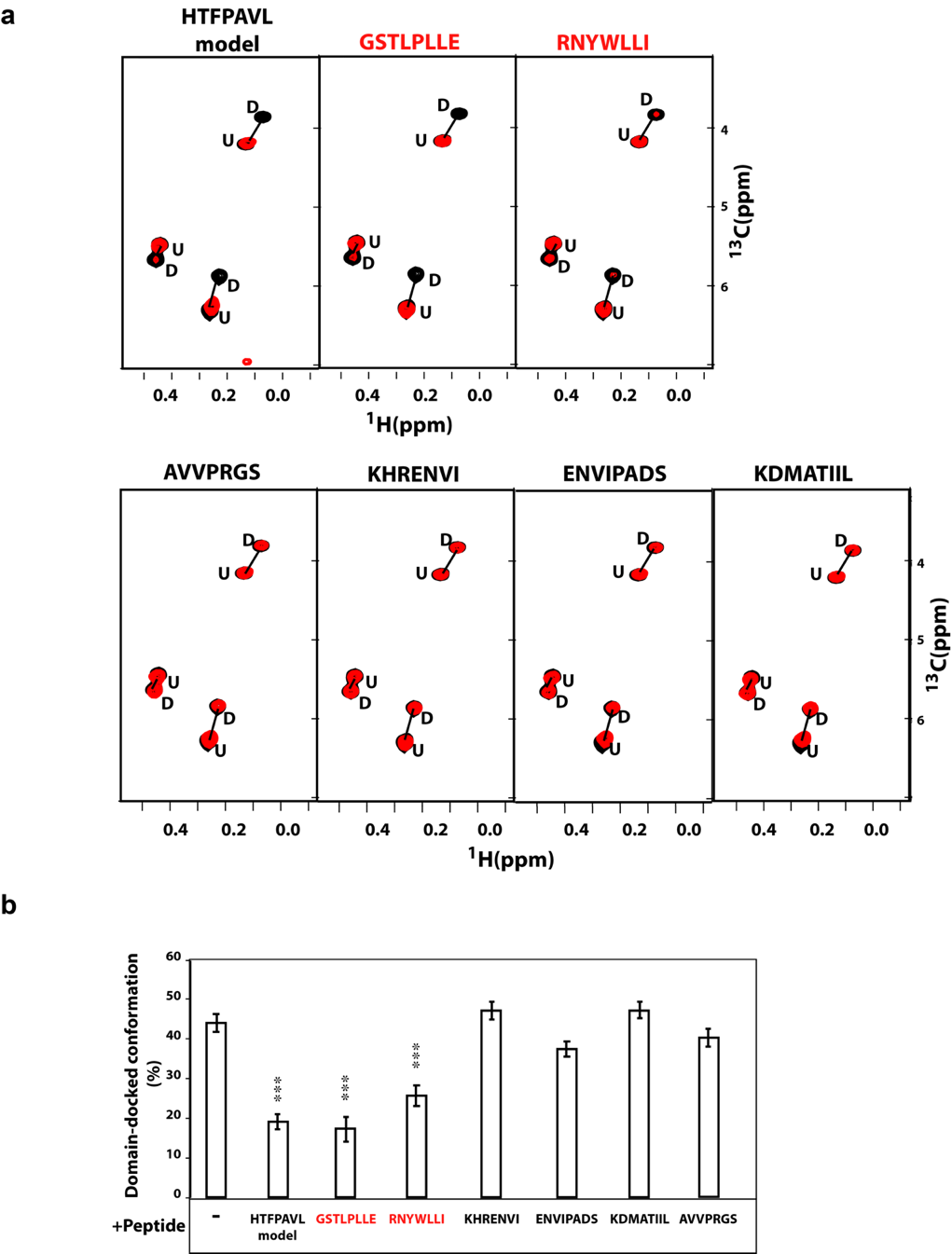

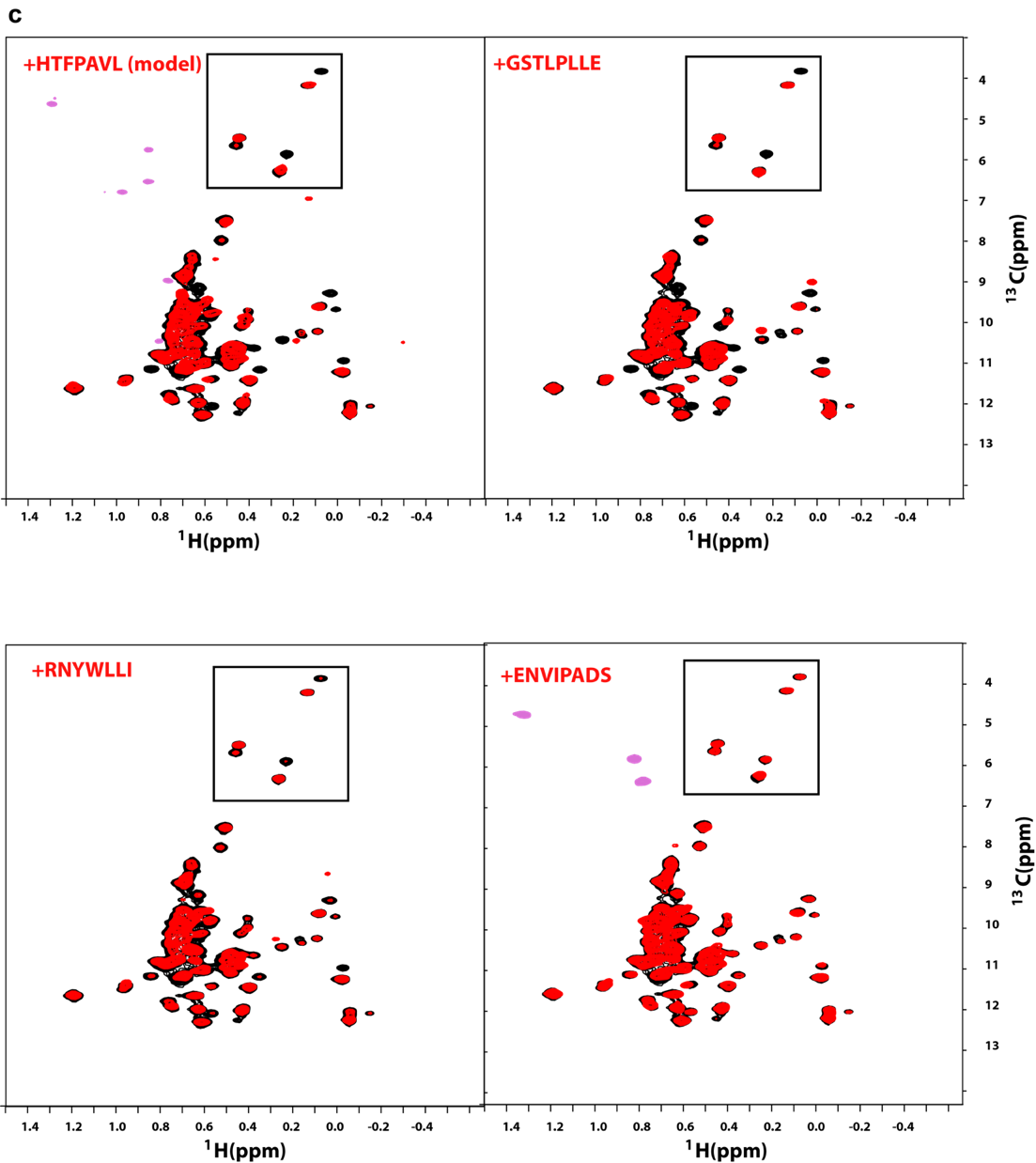

**Supplementary Figure 12: Perturbations in either BiP binding motif are insufficient to prevent substrate-induced oligomerization and BiP-dependent de-oligomerization.**

The SDS-PAGE analysis of de-oligomerization of WT Ire1-LD and its ^315^LL^316^ to DA and ^359^WLLI^362^ to GSSG variants (annotated) in the presence and the absence of BiP and ATP. 30 µM Ire1-LD were incubated 400 µM ΔEspP for three hours; if required 3 µM BiP and 40 mM ATP were added to the reaction and incubated for another 3 hours, following by collecting soluble and insoluble fractions. The addition of 400 µM ΔEspP results in formation of insoluble oligomers for WT Ire1-LD and the variants, which partially de-oligomerized in the presence of BiP and ATP.

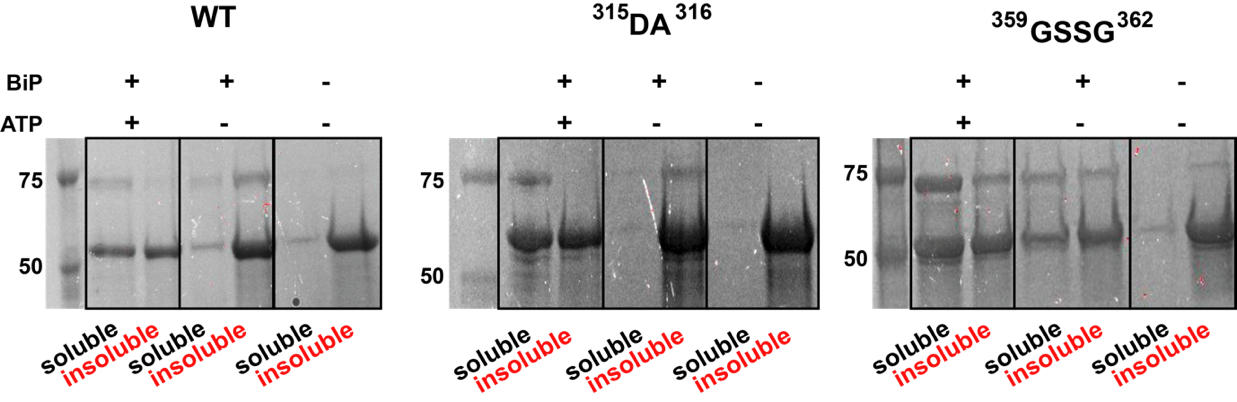

**Supplementary Figure 13: The sequence conservation of Ire1-LD.**

(a) Consurf^5,6^ conservation scores across the Ire1-LD amino acid sequence. The human Ire1 sequence (UniProt ID O75460) was used as an input sequence for the analysis, resulting in 124 HMMER-identified homologues with a minimal percentage identity between homologues of 30%.

(b) Simplified circular graph of the phylogenetic tree constructed by the Consurf^5,6^ and visualized by iTOL^7^; Ire1α or Ire1β sub-family members^8^ are highlighted by red and blue colours, respectively.

**a**

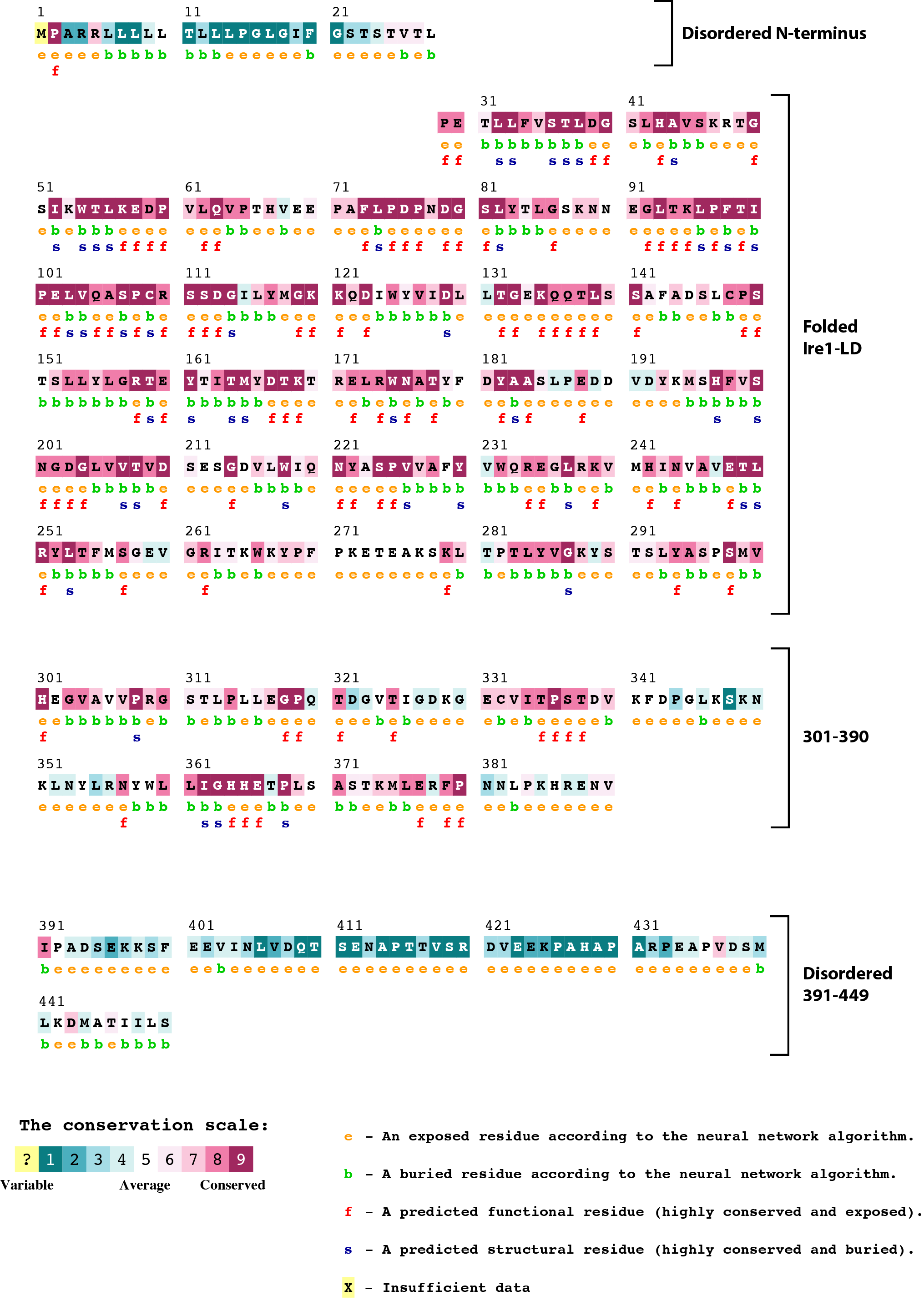

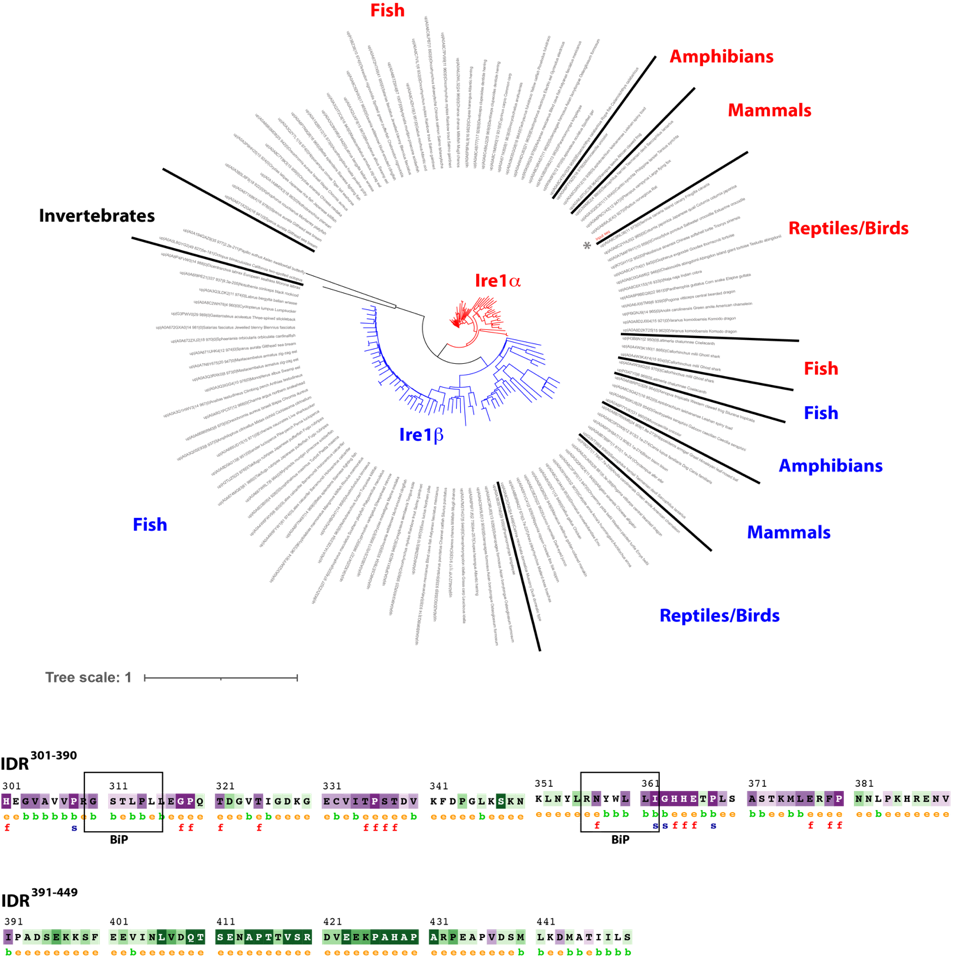

**b**

**Supplementary Figure 14: The effect of C-terminal truncations on Ire1-LD expression and solubility.**

The SDS-PAGE analysis of the *E. coli* expression of WT Ire1-LD (residues 24 to 449, left) and its truncated variants. The proteins were expressed at 37^o^C (a) and 20^o^C (b). For the variant that consists of residues 24 to 390 (middle), the entire juxtamembrane region (residues 391 to 449) was truncated. The variant that consists of residues 24-356 (right) also lacks the significant part of the oligomerization interface, including the highly conserved ^363^GHH^365^ β-strand. The whole cell *E. coli* lysate, along with its soluble and insoluble fractions, are shown. Truncation of the juxtamembrane region led to a slight increase in the amount of the insoluble protein when the protein was expressed at 37^o^C. However, at lower temperature (at 20^o^C), the 24-390 variant is predominantly soluble. In contrast, the truncation of the ^363^GHH^365^ β-strand significantly impacted Ire1-LD solubility at both temperatures.

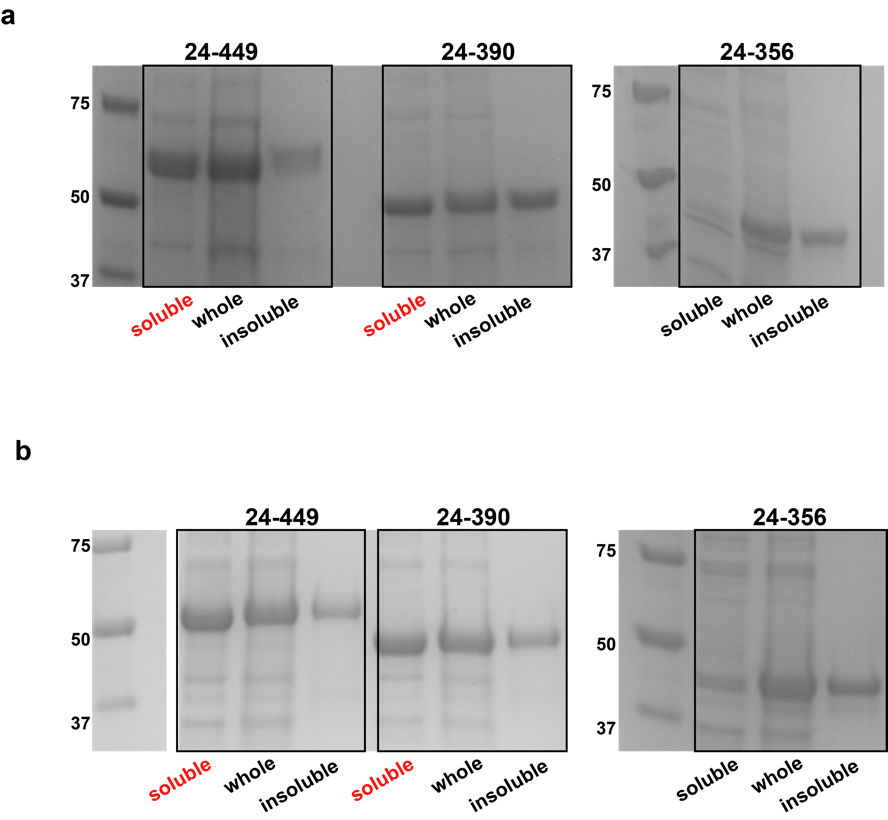

**Supplementary Figure 15: Dynamic properties of the Ire1-LD C-terminal region.**

(a) The amide 2D spectrum of WT Ire1-LD (black) overlaid with the spectrum of the core Ire1-LD (residues 24-390) that lacks the C-terminal region^391-449^ but contains BiP-binding motifs (green). For the core Ire1-LD construct, only 24 amide peaks (from ca. 300 and 90 expected for the folded Ire1-LD and flexible region 301-390) were visible in the 2D spectrum. In full agreement with previous observations^3^, the lack of peaks for the majority of core Ire1-LD and low peak intensities of the visible 24 peaks (also in b) suggest that the entire core region, including residues 301-390, is affected by μs-ms conformational dynamics. Consequently, the flexible region comprising residues 301-390 is predominantly ‘NMR invisible’, suggesting direct communication between residues 301-390 and the folded part of Ire1-LD. In contrast, most residues from the region that comprises residues 391-449 are present in the NMR spectrum (57 peaks from 59 expected), and the peaks from this region have significantly higher intensities (also in b), suggesting that residues 391-449 are not significantly affected by the μs-ms process in the core and, thus, do not directly communicate with the folded part of Ire1-LD.

(b) Peak intensities (calculated as a corresponding peak height divided by the noise level) for individual peaks in the amide 2D spectrum of WT Ire1-LD (a, black). The visible peaks were separated into two groups: (i) peak from flexible regions located in the core Ire1-LD (green) and (ii) peaks from the flexible region comprising residues 391-449 (black). The majority (>85%) of peaks for residues 391-449 has peak intensities larger than 10 (S/N), while the majority (>80%) of peaks from disordered core regions (residues <390) has peak intensity smaller than 10 (S/N). Significantly higher peak intensities for the residues 391-449 suggest that this region is largely independent from folded Ire1-LD and, thus, significantly less affected by its μs-ms conformational dynamics.

(c) Amide temperature gradients (ppb/K) calculated from chemical shifts obtained from the amide spectra of WT Ire1-LD recorded at 5, 10, 15, 20, and 25, ^o^C using linear regression. All residues with temperature gradients that occurs within the threshold of -3.6ppb/K were considered to adopt no secondary structure^9^. The temperature gradients were plotted separately for the C-terminal region 391-449 and the core region (residues <390). With low proton dispersions observed in the amide spectrum (a), characteristic amide temperature gradients for these peaks indicate that only disordered regions of Ire1-LD are NMR visible, while the folded part of Ire1-LD is NMR invisible.

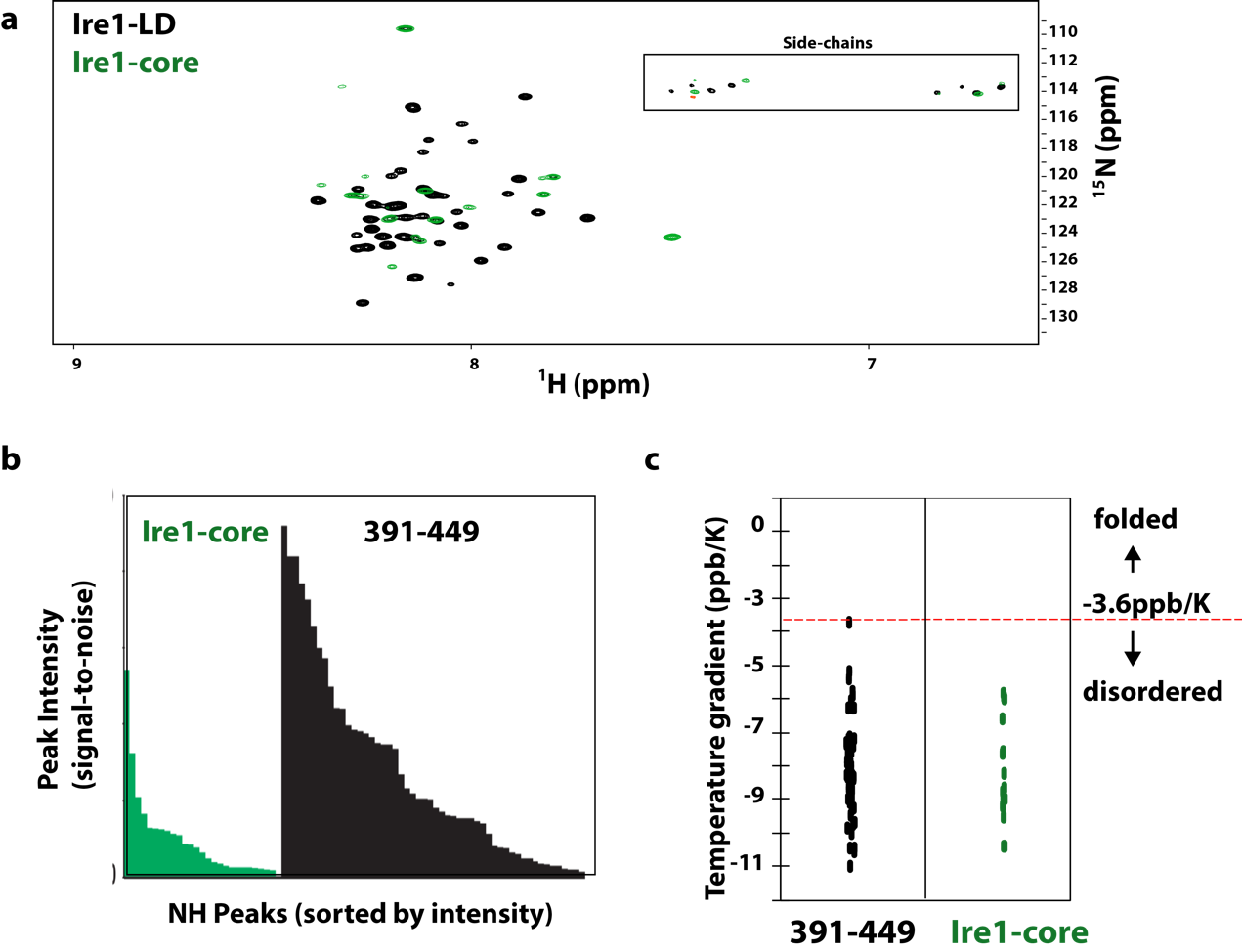

**Supplementary Figure 16: Conformational properties of the Ire1-LD C-terminal region.**

(Top) Secondary structure elements observed in the high-resolution X-ray structures of Ire1-LD (PDB IDs 2HZ6^1^ and 6SHC^10^). The ^306^VVP^308^ and ^363^GHH^365^ β-strands observed in both structures are shown in blue; while the α-helix adjacent to ^363^GHH^365^ (pink) is only formed in the 6SHC structure. The residues that unresolved in the X-ray structures are shown in yellow; grey shows the residues that were truncated from the X-ray constructs.

(Bottom) Predictions of disordered regions by several prediction algorithms, including IUPred^11^, ESpritz^12^, DISOPRED3^13^, MoreRONN^14^, PONDR^15^ and MobiDP lite^16,17^. Predicted regions of disorder are shown in yellow; the rest is coloured in white. The analysis revealed that the juxtamembrane region (residues 390 to 449) has classical characteristics of an IDR as the majority of this region is predicted to be disordered by all algorithms. In contrast, all algorithms failed to predict conformational disorder for most residues 301 to 390.

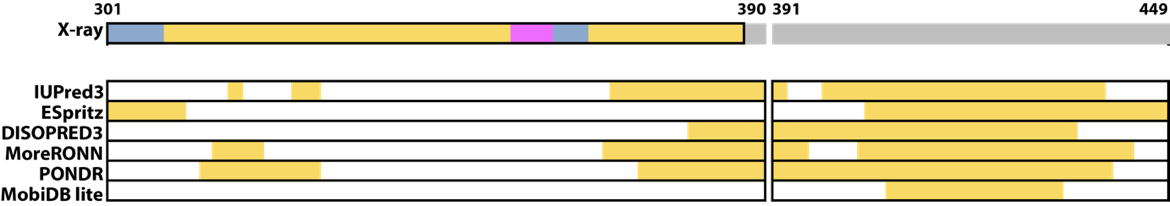

**Supplementary Figure 17: BiP solubilises insoluble Ire1-LD oligomer in the presence of 1-40 mM ATP.**

(A) The SDS-PAGE analysis of de-oligomerization of WT Ire1-LD in the presence and absence of BiP and different concentrations of ATP. 20 µM Ire1-LD was incubated 0 and 200 µM ΔEspP for 3 hours. If required 2 µM BiP and 0, 1, 10, or 40 mM ATP were added to the reaction and incubated for another 30 minutes, followed by collecting soluble and insoluble fractions. 10 µM lysozyme (band around 14 kDa) was added to each reaction as a loading control.

(B) The fraction of soluble Ire1-LD (same conditions as for A) monitored by the solubility assay.

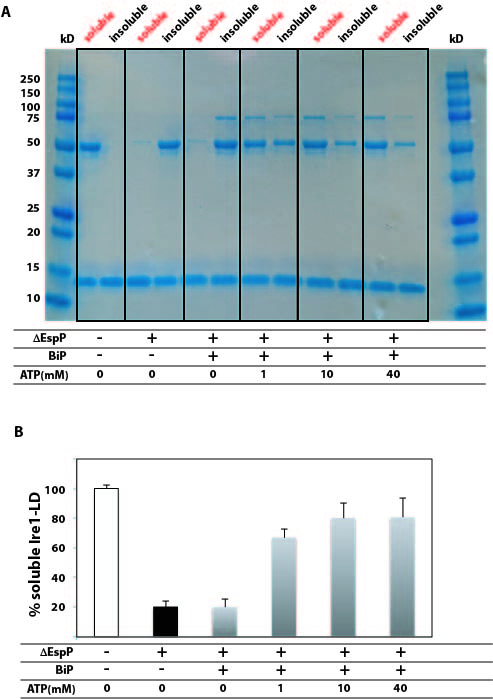

**Supplementary Table 1: Ire1-LD derived peptides predicted to be BiP binders by the BiPPred algorithm**

Predicted BiPPred^18^ scores of six motifs located in the C-terminal part of Ire1-LD. NMR experiments (Figure 3 and Supplementary Figure 11) confirmed BiP binding to two of these motifs (highlighted in yellow).

| **Sequence motif** | **Forward Score** | **Backward Score** | **Max Score** | **Confirmed (NMR)** |
| --- | --- | --- | --- | --- |
| **AVVPRGS**  ^305^AVVPRGS^311^ | 0.9517 | 0.6762 | 0.9517 | **No** |
| **GSTLPLLE**  ^310^GSTLPLL^316^  ^311^STLPLLE^317^ | 0.3335  0.949 | 0.8632  0.9509 | 0.8632  0.9509 | **Yes** |
| **RNYWLLI**  ^356^RNYWLLI^362^ | 0.9427 | 0.943 | 0.943 | **Yes** |
| **KHRENVI**  ^385^KHRENVI^391^ | 0.3792 | 0.8463 | 0.8463 | **No** |
| **ENVIPADS**  ^388^ENVIPAD^394^  ^389^NVIPADS^395^ | 0.8071  0.8287 | 0.7215  0.2751 | 0.8071  0.8287 | **No** |
| **KDMATIIL**  ^442^KDMATII^448^  ^443^DMATIIL^449^ | 0.5798  0.3383 | 0.7999  0.9265 | 0.7999  0.9265 | **No** |

**Supplementary Table 2: Variations in the sequences of the BiP binding motifs does not significantly affect their ability to bind BiP**

Predicted BiPPred scores^18^ for the ^310^GSTLPLL^316^ and ^356^RNYWLLI^362^ BiP binding motifs calculated for several representative Ire1 sequences (organisms and UniProt IDs are annotated). Characteristic amino acid variations in these motifs among different Ire1s are shown in red. These variations do not significantly affect the ability of these motifs to bind BiP, as suggested by the fact that the majority of sequences have a BiPPred score of 0.84 and higher (very good binders, dark yellow), and few sequences have a score of 0.7-0.84 (good binders, light yellow).

| Organism | UniProt ID | Sequence motif | Forward score | Backward score | Max score |
| --- | --- | --- | --- | --- | --- |
| **^310^GSTLPLL^316^** | | | | | |
| **Ire1α** | | | | | |
| Human | O75460 | GSTLPLL | 0.3335 | 0.8632 | 0.8632 |
| Tortoise | A0A8C4Y7H0 | GRAIPLL | 0.599 | 0.9006 | 0.9006 |
| Bird | A0A8C9NL08 | GSAIPLL | 0.2908 | 0.9152 | 0.9152 |
| Frog | A0A8J0TUC3 | GRAIPLL | 0.599 | 0.9006 | 0.9006 |
| Fish | A0A3B3RS82 | GSTFPLL | 0.3335 | 0.8632 | 0.8632 |
| Fish | A0A4W4DU83 | GSTFPML | 0.3528 | 0.7572 | 0.7572 |
| **Ire1β** | | | | | |
| Human | Q76MJ5 | GLTLAPA | 0.8926 | 0.5888 | 0.8926 |
| Birth | A0A8C4K344 | GITLARI | 0.9279 | 0.757 | 0.9279 |
| Turtle | A0A8C3F3F0 | GITLARI | 0.9279 | 0.757 | 0.9279 |
| Toad | A0A8C5Q421 | GITLAQV | 0.9441 | 0.8477 | 0.9441 |
| Fish | A0A6P7N497 | GLTLARI | 0.9152 | 0.7596 | 0.9152 |

| **^356^RNYWLLI^362^** | | | | | |
| --- | --- | --- | --- | --- | --- |
| Human | O75460 | RNYWLLI | 0.9427 | 0.943 | 0.943 |
| Tortoise | A0A8C0GAW6 | RNHWLLI | 0.8083 | 0.6218 | 0.8083 |
| Alligator | A0A3Q0H0Z4 | HNQWLLI | 0.9346 | 0.9172 | 0.9346 |
| Toad | A0A8C5R512 | RNQWLLI | 0.9346 | 0.9172 | 0.9346 |
| Fish | A0A3Q3JXF9 | RNYLLLI | 0.9427 | 0.943 | 0.943 |
| Fish | A0A8C4B6J2 | KNHLLLI | 0.8083 | 0.6218 | 0.8083 |
| Fish | A0A6J2VYF1 | QNQWLLI | 0.9346 | 0.9172 | 0.9346 |

**References:**

1.	Zhou, J. et al. The crystal structure of human IRE1 luminal domain reveals a conserved dimerization interface required for activation of the unfolded protein response. *Proc Natl Acad Sci U S A* **103**, 14343-8 (2006).

2.	Borgia, M.B., Nickson, A.A., Clarke, J. & Hounslow, M.J. A mechanistic model for amorphous protein aggregation of immunoglobulin-like domains. *J Am Chem Soc* **135**, 6456-64 (2013).

3.	Karagoz, G.E. et al. An unfolded protein-induced conformational switch activates mammalian IRE1. *Elife* **6**(2017).

4.	Wieteska, L., Shahidi, S. & Zhuravleva, A. Allosteric fine-tuning of the conformational equilibrium poises the chaperone BiP for post-translational regulation. *Elife* **6**(2017).

5.	Ashkenazy, H. et al. ConSurf 2016: an improved methodology to estimate and visualize evolutionary conservation in macromolecules. *Nucleic Acids Res* **44**, W344-50 (2016).

6.	Ben Chorin, A. et al. ConSurf-DB: An accessible repository for the evolutionary conservation patterns of the majority of PDB proteins. *Protein Sci* **29**, 258-267 (2020).

7.	Letunic, I. & Bork, P. Interactive Tree Of Life (iTOL) v5: an online tool for phylogenetic tree display and annotation. *Nucleic Acids Res* **49**, W293-W296 (2021).

8.	Cloots, E. et al. Evolution and function of the epithelial cell-specific ER stress sensor IRE1beta. *Mucosal Immunol* **14**, 1235-1246 (2021).

9.	Cierpicki, T. & Otlewski, J. Amide proton temperature coefficients as hydrogen bond indicators in proteins. *J Biomol NMR* **21**, 249-61 (2001).

10.	Amin-Wetzel, N., Neidhardt, L., Yan, Y., Mayer, M.P. & Ron, D. Unstructured regions in IRE1alpha specify BiP-mediated destabilisation of the luminal domain dimer and repression of the UPR. *Elife* **8**(2019).

11.	Erdos, G., Pajkos, M. & Dosztanyi, Z. IUPred3: prediction of protein disorder enhanced with unambiguous experimental annotation and visualization of evolutionary conservation. *Nucleic Acids Res* **49**, W297-W303 (2021).

12.	Walsh, I., Martin, A.J., Di Domenico, T. & Tosatto, S.C. ESpritz: accurate and fast prediction of protein disorder. *Bioinformatics* **28**, 503-9 (2012).

13.	Jones, D.T. & Cozzetto, D. DISOPRED3: precise disordered region predictions with annotated protein-binding activity. *Bioinformatics* **31**, 857-63 (2015).

14.	Ramaraj, V. Oxford University (2014).

15.	Xue, B., Dunbrack, R.L., Williams, R.W., Dunker, A.K. & Uversky, V.N. PONDR-FIT: a meta-predictor of intrinsically disordered amino acids. *Biochim Biophys Acta* **1804**, 996-1010 (2010).

16.	Piovesan, D. et al. MobiDB: intrinsically disordered proteins in 2021. *Nucleic Acids Res* **49**, D361-D367 (2021).

17.	Potenza, E., Di Domenico, T., Walsh, I. & Tosatto, S.C. MobiDB 2.0: an improved database of intrinsically disordered and mobile proteins. *Nucleic Acids Res* **43**, D315-20 (2015).

18.	Schneider, M. et al. BiPPred: Combined sequence- and structure-based prediction of peptide binding to the Hsp70 chaperone BiP. *Proteins* **84**, 1390-407 (2016).
